# Learning Environmental Dynamics: Building Internal Models of a Simulated External Force

**DOI:** 10.64898/2026.09.17.751840

**Authors:** Jacob J. Boulrice, Bernard Marius ‘t Hart, Denise Y. P. Henriques

## Abstract

Motor adaptation is typically studied using limb-based perturbations in highly constrained tasks. However, everyday actions require compensating for uncertain environmental dynamics impacting manipulated objects within redundant execution spaces, where multiple combinations of variables can achieve success. To investigate this, we designed a novel virtual reality task where participants launched a ball across a lateral water current to spatially distinct targets. Two experiments (2.0 m/s and 3.0 m/s) tested if classic adaptation signatures remain consistent across varying perturbation sizes.

Results revealed three key signatures. First, participants exhibited robust error reduction to all targets during training. Second, between-subjects analyses revealed that this learning produced substantial, immediate transfer to untrained target locations. Finally, persistent motor aftereffects appeared upon perturbation removal, despite explicit cues indicating absence of the perturbation. Together, these results demonstrate that humans form internal models under altered projectile–environment dynamics, and that prior experience facilitates broad generalization across the workspace.

## Introduction

We live in a dynamic world where external forces such as gravity, wind, and water currents constantly shape the relationship between our actions and their outcomes. To maintain accurate control, the central nervous system (CNS) must predict how the body, manipulated objects, and surrounding environment will influence action outcomes and update these predictions when errors occur. To account for these changing conditions, the CNS builds and continuously recalibrates internal models—neural representations that predict how motor commands, body mechanics, manipulated objects, and the environment interact to estimate the sensory consequences of an action. During movement execution, internal forward models use an efference copy of the motor command—a neural representation of the intended movement—to predict future sensory consequences of the motor command (Wolpert et al., 1995). When a discrepancy arises between predicted and observed sensory feedback, a sensory prediction error (SPE) occurs, which serves as the primary signal to update these internal models (Tseng et al., 2007). For example, a golfer competing in a tournament may encounter several changes that require adaptation. Fatigue changes the properties of the body, requiring adjustments to motor commands to maintain performance. Switching to unfamiliar clubs changes the dynamics of the tool, altering how forces are transmitted to the ball. Finally, changing environmental conditions, such as strong wind, modifies the ball’s flight after it has been struck. Although each of these sources of change facilitate and require adaptation, they differ in what must be predicted or estimated. Changes to the body primarily require updating predictions about one’s own movements, changes to tools require learning how the tool transforms those movements, whereas environmental forces require predicting how external dynamics will influence an object’s motion after the action has been completed. Importantly, these sources of change differ in the degree or quality of information available to the CNS. When adapting to changes in the body, the CNS receives relatively direct proprioceptive and visual feedback about the limb position and movement. Similarly, the properties of tools may be partially observable through haptic and visual signals. In contrast, environmental dynamics such as wind or water currents primarily influence the manipulated object after it leaves the body or tool. Consequently, the CNS receives little or no proprioceptive information about how these external forces alter the object’s motion. Instead, the environmental dynamics must be inferred from contextual cues, such as visible water flow or wind, together with observed consequences of previous actions (Berniker & Körding, 2008; Shadmehr & Mussa-Ivaldi, 1994). Consequently, learning environmental dynamics may place unique demands on prediction and motor adaptation compared to traditional, limb-based adaptation paradigms.

Despite the importance of adaptation to changes in the body, tool, and environment, much of our understanding of sensorimotor adaptation comes from experimental paradigms that primarily perturb limb movements directly, providing comparatively less insight into how humans adapt to altered environmental dynamics on manipulated objects. Our understanding of human sensorimotor learning and adaptation is therefore largely based on controlled experimental paradigms such as visuomotor rotations (VMRs), visuomotor gains, and force fields (FFs).

These visuomotor and force field adaptation paradigms have shown that the human CNS updates internal models in response to SPEs (Shadmehr & Mussa-Ivaldi, 1994; Smith et al., 2006; Tseng et al., 2007). Learning in these paradigms is retained through context-dependent processes (Heald et al., 2021), generalizes to novel contexts as a function of similarity to the training context (Heuer & Sülzenbrück, 2013; Thoroughman & Shadmehr, 2000), and shows persistent compensatory movements (i.e, motor aftereffects) upon sudden removal of the perturbation (Shadmehr & Mussa-Ivaldi, 1994). Furthermore, motor learning often reflects interaction between explicit adjustments—conscious, goal-directed, and strategic movements due to task errors—and implicit recalibration, an unconscious process that gradually updates internal model expectations in response to SPEs (Morehead et al., 2017; Tsay et al., 2022; Wilterson & Taylor, 2021).

While powerful and experimentally tractable, these visuomotor and force field paradigms constrain adaptation to relatively low-dimensional solution manifolds. Here we use the term solution manifold to refer to the set of execution variable combinations that successfully achieve the task goal. Some motor learning tasks have only a narrow range of successful movement solutions, whereas others allow many different combinations of execution variables to achieve the same outcome. For example, in a classic VMR task, participants move toward a visual target (red circle in Figure 1), but the visual feedback representing their movement direction is experimentally rotated relative to the true movement direction. As illustrated in Figure 1A, while the actual hand movement is directed straight toward the target (dashed line), the visual cursor feedback is displaced by an imposed rotation (black arrow; 30° clockwise in the example). With practice, participants gradually alter their unseen hand movements to compensate for this discrepancy, adjusting movement direction such that the rotated cursor lands on the target as shown in Figure 1B. Because successful performance depends primarily on modifying a single execution variable—movement direction—the available solution manifold is effectively one-dimensional; similar to the solution manifold depicted in Fig 3A, but with the direction shifted 30° to the left/counter-clockwise). Thus, adaptation largely involves recalibrating the mapping between hand movement and visual cursor feedback while leaving the underlying task structure unchanged.

**Figure 1:**
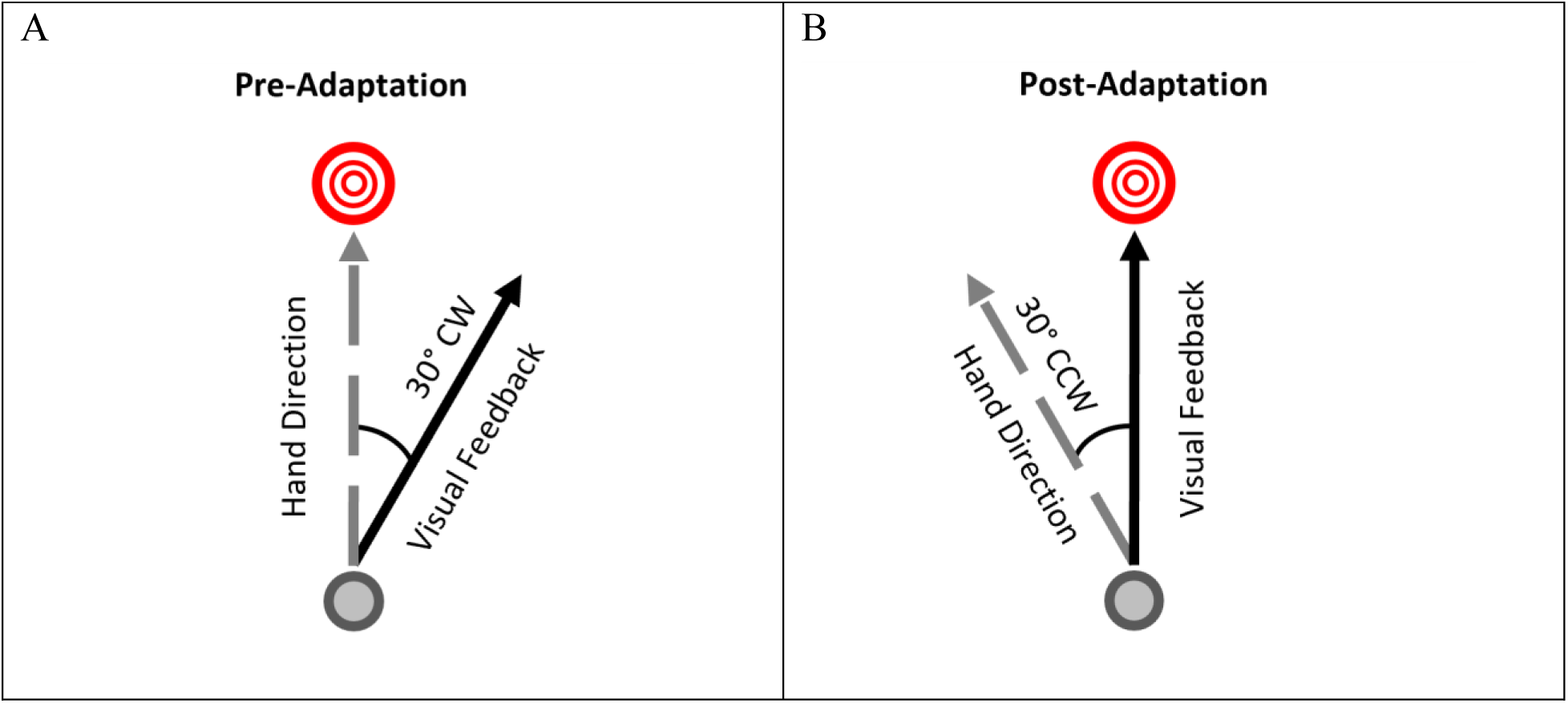
Compensating for One-Dimensional Visuomotor Rotations (VMRs). VMRs represent a substantial amount of our knowledge surrounding motor learning and adaptation. In the above example, the participant must move the cursor (lower gray circle) to the upper red target. A: If their movement direction follows the gray dashed arrow while a 30° clockwise (CW) rotation is active, the cursor visual feedback will move along the trajectory of the solid black arrow. B: Adaptation in this case is the process of adjusting one’s movement direction toward the 30° counterclockwise (CCW) gray dashed arrow. Here, the solution manifold is one-dimensional such that a single execution variable (i.e., movement direction) dictates task success. This form of adaptation involves adjusting visual expectations associated with reaches (i.e., adapting the hand-cursor mapping). While this manipulation has been well-studied and characterizes adaptation with robust experimental control, the ecological validity of such a perturbation is debatable given humans rarely encounter them in the real world.

Although traditional adaptation paradigms have provided substantial insight into sensorimotor learning, they often preserve relatively similar task structures between perturbed and unperturbed conditions, differing primarily in the movement direction required for success. Moreover, perturbations in these paradigms are typically inferred from changes in movement outcome rather than from directly observing the perturbation itself. In contrast, real-world environments provide continuous visual and auditory information about environmental conditions (e.g., visible wind or water flow), while still requiring individuals to predict how these conditions will influence object motion.

Moreover, while previous work using VMRs and FFs has substantially advanced our understanding of sensorimotor learning and adaptation in response to direct manipulations of limb movements, it remains less clear how the CNS adapts to environmental perturbations that act externally on manipulated objects. Unlike traditional paradigms, where perturbations directly alter the relationship between limb movement and sensory feedback, environmental perturbations require individuals to infer and predict the dynamics of external forces that alter object motion after an action has already been executed. Consequently, it remains unclear whether key behavioural signatures commonly observed in traditional paradigms—including error reduction, transfer, and aftereffects—extends to environmental perturbations.

Previous work by Pouke et al., (2024) provided evidence that the CNS can adapt to altered object dynamics in a goal-directed virtual reality (VR) ball tossing task under simulated hypergravity (5g). Importantly, when participants returned to the normal gravity condition, participants exhibited compensatory aftereffects that gradually decayed despite explicit awareness that gravity had returned to baseline levels. These findings suggest that the CNS can modify internal models of object-environment dynamics and retain these adaptations even after the perturbation is removed.

The present study aims to extend this work by introducing a novel VR framework to investigate motor adaptation to a simulated water current perturbation. This thesis builds upon the work of Modchalingam et al. (2025), where participants adapted to the effects of simulated gravity while rolling a virtual ball across a visually slanted surface in VR. While the geometric tilt of the surface was visible and informative in their paradigm, gravity itself is not directly observable.

Therefore, its magnitude and influence on the ball’s trajectory had to be inferred indirectly from post-movement sensory feedback and the geometric properties of the immediate environment. In contrast, the present framework may provide richer cues, by introducing a simulated water current where the physical perturbation is perfectly coupled with the visual state of the water.

As illustrated in Figure 2 (and demonstrated in the video provided in the QR code), participants launched a virtual ball toward one of four spatially distinct targets positioned across a flat-water surface while compensating for a leftward water current perturbation. The visual motion of the water surface and accompanying flowing-water audio provided continuous cues regarding the presence of the perturbation; potentially reducing ambiguity regarding the environmental state. Unlike traditional perturbations that directly alter the relationship between limb movements and sensory feedback, the present perturbation acts only on the ball after release. Consequently, participants must predict how the environment will alter object motion, rather than compensate for changes in the movement of their own limb.

**Figure 2:**
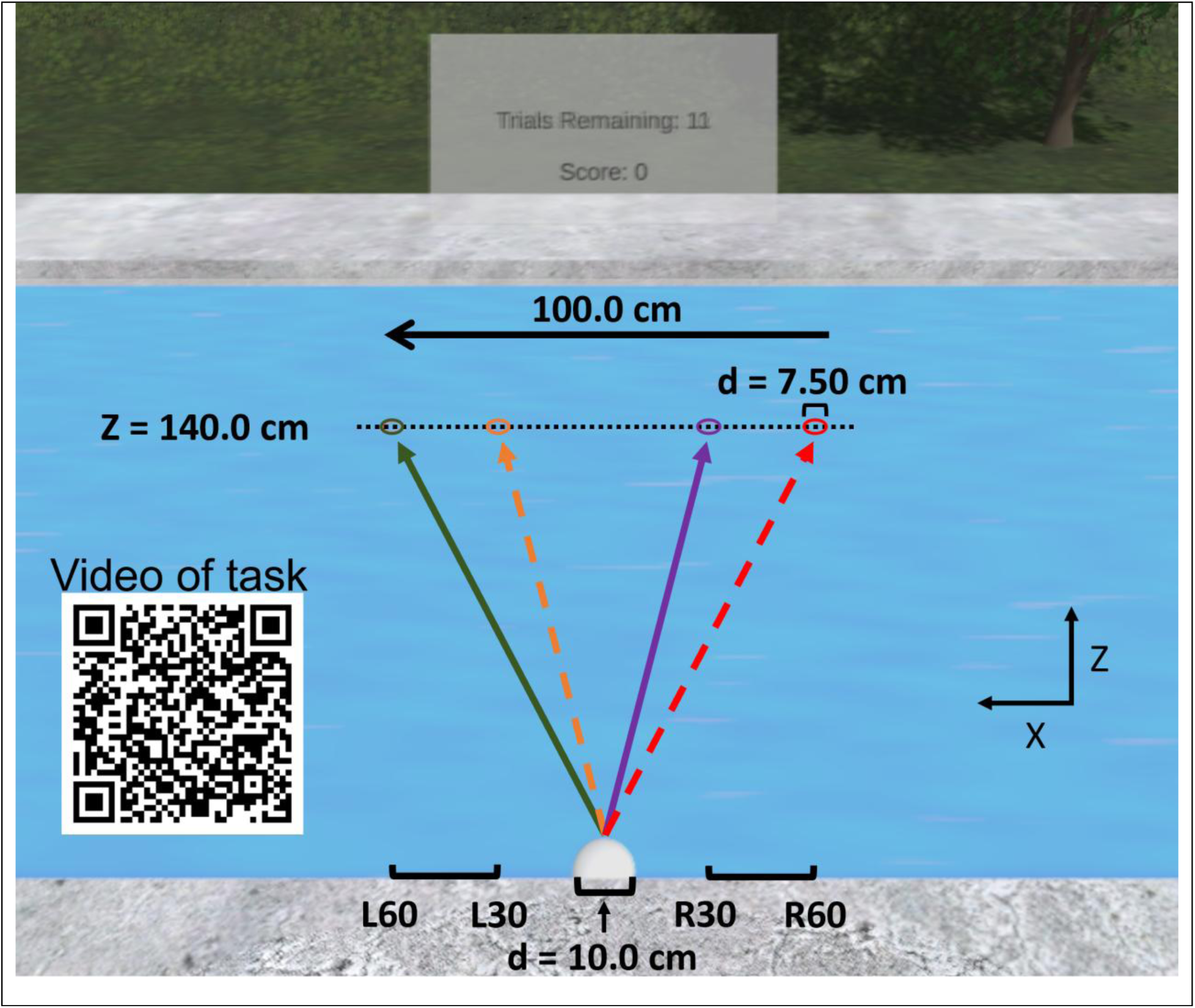
Overview of the Virtual Ball-Launching Task and Environmental Context. When the perturbation is active, the water current pushes the ball leftward (downstream) at either 2.0 or 3.0 m/s (depending on the condition). Arrow line-styles indicate target pairs, where participants were exposed to one downstream (L: leftward) and one upstream (R: rightward) target per Training and Washout phase. For example, one group experienced the solid line targets (L60 and R30) in the Training Phase and switched to the dashed targets (L30 and R60) in the Transfer Phase, while the other group experienced the opposite order. Line-styles and target colours are for visualization purposes only; during a trial, participants did not see the arrows and only saw a single red target ring.

Importantly, adaptation in the present task differs from traditional paradigms in another fundamental way: successful performance emerges within a redundant solution manifold–that is, multiple combinations of launch directions and speeds can achieve the same task goal. Previous work on motor learning in multidimensional solution manifolds suggests that learners initially exploit broad regions of low error before refining behaviour toward motor-equivalent solutions that preserve task success while accommodating intrinsic motor variability (Zhang et al., 2018; Zhang & Sternad, 2021). However, unlike paradigms such as the Skittles task conducted in Zhang et al. (2018)—where multi-dimensional solution manifolds are intrinsic to task mechanics—the present study introduces a clear environmentally specific perturbation that fundamentally transforms the task landscape. As illustrated in Figure 3, the unperturbed environment affords a relatively constrained, one-dimensional solution manifold in which movement direction is the primary determinant of success (Figure 3A). In contrast, the addition of a lateral water current dynamically reshapes the task into a redundant solution manifold, where successful performance can be achieved through multiple combinations of launch deviation and speed (Figure 3B). Thus, participants may compensate for the same perturbation through distinct movement solutions that achieve the same outcome, introducing greater flexibility in action selection. Here, learning is not a simple remapping of hand direction driven by sensory prediction errors. Rather, it’s a process of navigating the execution space to discover successful combinations of launch direction and speed that lie on the solution manifold.

**Figure 3:**
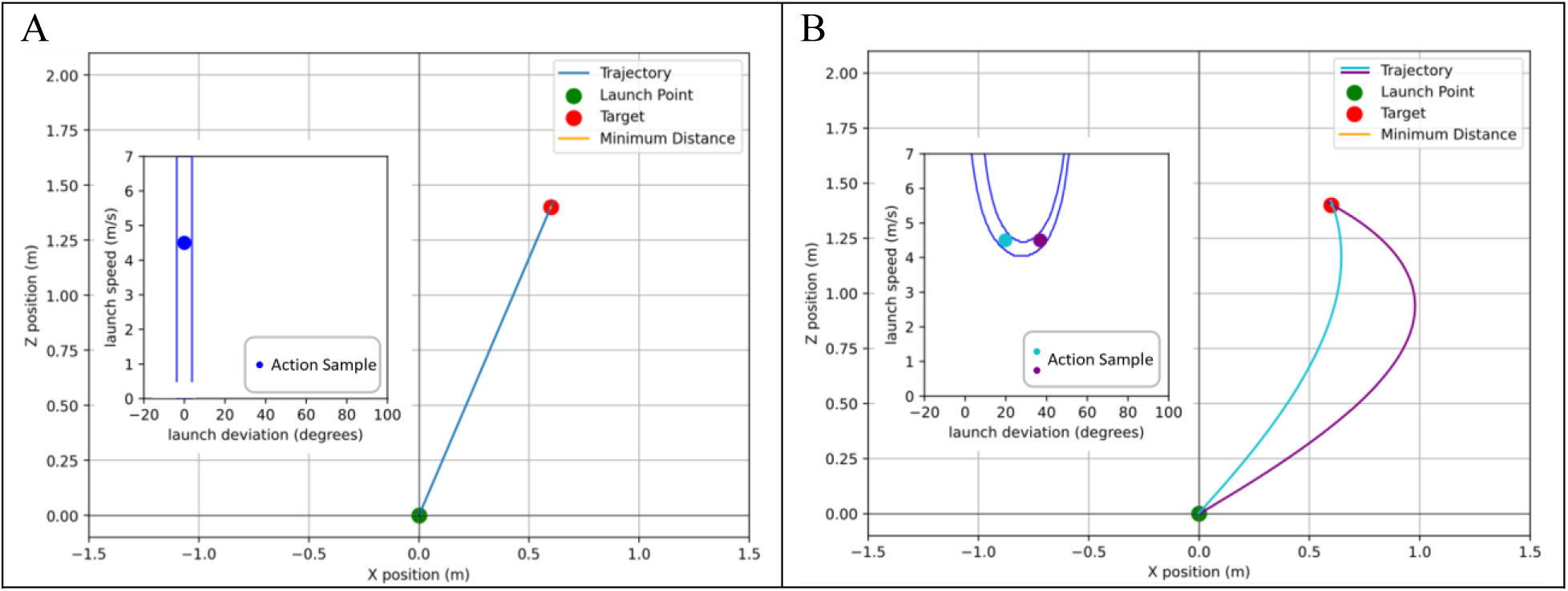
Comparing Non-Redundant and Redundant Solution Manifolds. A task is considered redundant when an infinite set of task-relevant execution variables can be executed to satisfy the goal (Zhang et al., 2018). The above figures illustrate simulations of this thesis’ present task, where participants completed a ballistic ball launching task under dynamic environmental conditions. Participants modulated two relevant to counter the lateral perturbation: launch deviation and speed. Both figures illustrate trajectories from launches at 4.5 m/s. A: unperturbed trajectory illustrating a one-dimensional solution manifold, requiring a launch deviation of 0.0° to intersect with the target centre, while all speeds above threshold are successful. B: Perturbed trajectories with the same launch speed (4.5 m/s) can intersect the target centre at two distinct launch deviations relative to target (19.8° and 36.8°) to the right, demonstrating a redundant solution manifold.

Understanding how the CNS learns and generalizes under environmentally specified perturbations with multi-dimensional solution manifolds remains an open question. The present study therefore aimed to examine motor learning and adaptation to dynamic environmental perturbations during a redundant virtual ball-launching task.

Here, we test whether humans form predictive action-outcome mappings of object-environment dynamics, if this learning generalizes across target locations within the same environment, and whether adaptation produces aftereffects following sudden removal of the perturbation. We hypothesize that participants would gradually reduce performance error during training, that environmental state would act as contextual cues for supporting generalization to novel target locations, and that learning would produce modest, but measurable aftereffects following perturbation removal. More broadly, the present paradigm is intended to provide a flexible and experimentally tractable framework for investigating how humans adapt to environmentally specified perturbations, offering new insight into the mechanisms underlying motor learning, generalization, and action selection in redundant, multi-dimensional environments.

## Methods

### Participants

A total of 83 participants (59 females, mean age = 20.71, SD age = 4.43) completed the study in one of two different waters current experiments (n = 38, 2.0 m/s experiment; n = 45, 3.0 m/s experiment). All participants were right-handed and had normal or corrected-to-normal vision. Participation was voluntary, and all participants gave informed, written consent prior to data collection. The procedures used in this study were approved by the York Human Participants Review Sub-committee. All participants were recruited through the Undergraduate Research Participant Portal (URPP) and through word of mouth. On completion of the study, URPP participants were compensated with course credits.

### Apparatus and VR Environment

Participants sat on a height-adjustable chair, positioned in front of a table. During the instruction period, they were given the details of the task environment and goal. Following the instructions, participants donned an Oculus Quest Pro head-mounted-display (resolution: 1800 by 1920 per eye; refresh rate of 72 Hz). The task was performed with the right-hand Quest Touch Pro controller (refresh rate of 72 Hz), which participants used to interact with the virtual environment. During the task, participants rested their forearm on the tabletop and initiated movements from a standardized home position located below the chin, near the body midline.

The immersive virtual environment was developed in Unity (version 2022.3.1f1) and depicted a flat-water surface extending in front of the participant. A virtual ball and target rings were displayed within the environment, and participants viewed the scene depicted in Figure 2 from a first-person perspective via the headset. Participants did not see a virtual representation of their hand or the controller to minimize additional visual information about limb position and encourage reliance on the observed ball trajectory. Depending on the experimental phase, the water surface was either stationary or visually moving to reflect the presence of a simulated water current. Unity’s fixed-timestep physics update (Δt = 0.01389 s, 72 Hz) was used to ensure consistent rendering and simulation of object dynamics.

### Task and Stimuli

Participants completed an immersive VR ball-launching task in which they attempted to launch a virtual ball toward visually presented target rings. Ball movement in the virtual environment was constrained to two dimensions such that its motion only occurred within the horizontal (x-axis; left-right) and frontal plane (z-axis; forward directions as illustrated in Figure 2). Restricting movement to these two dimensions isolated adaptation to the lateral water-current perturbation while reducing unnecessary task complexity. The ball’s path along the water was partially submerged to keep the physics of the water on the ball consistent across the entire trajectory, as demonstrated in the task video (see QR code in Figure 2).

On each trial, a single target ring (diameter = 7.5 cm) appeared in front of the participant, at one of four possible horizontal locations: 30.0 or 60.0 cm left and right of the midline positioned 140.0 cm in front of the participant (Figure 2). All targets were arranged along the same depth plane to be consistent with the direction of the lateral perturbation. This required participants to consistently execute launches that reached the target depth (Z = 140.0 cm), decelerating and flowing downstream along a shared plane that was congruent with the horizontal water current perturbation vector. Participants launched a virtual ball (diameter = 10.0 cm; virtual mass = 1.0 kg) from the home position toward the target with the goal of intersecting or passing as close as possible to it.

Participants initiated each throw from a standardized home position located below their chin—at the midline—and pressed the controller trigger to register the start position. To launch the ball, participants then performed a rapid forward arm movement, moving the controller 30.0 cm away from the start position or releasing the trigger. The direction and speed of this movement determined the initial launch velocity of the ball, allowing participants to vary both launch direction and launch speed to control the ball trajectory for task success. The initial launch velocity of the virtual ball was determined from the controller’s average velocity during the final 10 recorded samples immediately before release. A launch was registered when participants either released the trigger or moved the controller beyond the 30.0 cm displacement threshold, whichever occurred first. To initiate a trial, launch speed had to exceed 0.5 m/s. Slower movements were treated as unsuccessful launch attempts, and participants were prompted to throw faster before the trial was restarted. This threshold prevented weak or inadvertent movements from initiating a trial while ensuring that participants produced deliberate launch attempts.

### Experimental Design

Across the experiment, participants experienced two water-surface contexts: a no current (unperturbed) and an active (perturbed) current context. In the no-current context, the water surface was visually and functionally stationary, such that successful performance could only be achieved by launching the ball directly toward the target at any speed above threshold (i.e., a single effective movement solution or one-dimensional solution manifold). In the active-current context however, a constant leftward acceleration altered ball trajectories after launch, requiring participants to compensate by adjusting both launch direction and speed (i.e., a two-dimensional solution). Depending on experiment, the water current magnitude was either 2.0 m/s or 3.0 m/s. Given that this was a novel paradigm, we tested two water current magnitudes to see whether the primary signatures of adaptation would replicate under a perturbation that would produce larger errors and thus require greater behavioural compensation.

As depicted in Figure 4, both experiments progressed through 5 sequential phases: Baseline, Training Phase 1, Washout-Top-Up Phase 1, Training Phase 2, and Washout-Top-Up Phase 2. Each phase differed in the presence or absence of the moving water current, the frequency of water-state transitions, and the presented target pair. The target pair experienced during phases was counterbalanced across two groups, with group one training on L30 and R60, before workspace effectively transferred to novel targets locations, by comparing the initial performance for naive versus experienced participants on the same target.

**Figure 4:**
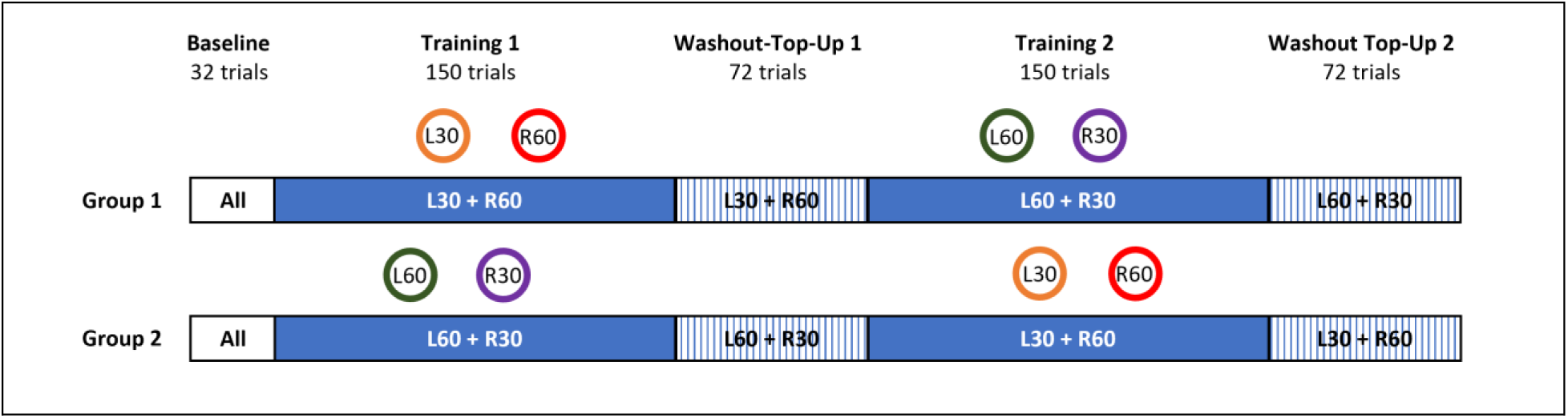
Experimental Trial Schedule. Participants initially complete 52 inactive-current trials for all four targets, followed by Training Phase 1 with 150 active-current trials, presenting a counterbalanced pair of targets (i.e., one left and right target). Next, during the Washout-Top-Up phase, the current was suddenly inactivated and activated every 8 trials with the same pair from the previous Training Phase, repeating a total of 72 trials. The Training Phases and Washout-Top-Up phases were repeated but with the opposite target pair to measure generalization, during the second Training and Washout-Top-Up Phases.

### Trial Schedule & Phases

#### *Familiarization and Baseline* [20 trials, 5 per target; 32 trials, 8 per Target]

Participants initially completed no-current trials with each of the four targets presented in a pseudorandomized order, with none appearing more than twice consecutively (applied in all phases). During Familiarization and Baseline, the water surface was visually and functionally stationary, first establishing familiarization with the ball-launching mechanics, and initial performance during Baseline.

#### *Training Phase 1* [150 trials, 75 per Target]

Immediately following Baseline, the leftward water-current perturbation was introduced (speed was either 2.0 m/s or 3.0 m/s depending on the experiment), which curved the ball trajectory into an arc relative to the launch direction. The presence of the current was indicated by visual motion of the water surface and the sound of flowing water. Here, participants practiced and learned to launch the ball to intersect a pair of spatially distinct targets (i.e., a left/downstream and right/upstream target). During this phase, half of the participants launched to targets L30 and R60, while the other half launched to targets L60 and R30 for their training target pair. This phase allowed us to observe initial learning of ball-environment dynamics in the absence of prior experience.

#### *Washout Top-up Phase 1* [72 trials, 36 per Target]

During the first Washout-Top-Up phase, the environment alternated between no-current trials and active-current trials in short cycles, presenting the training target pair (see Figure 4). Each cycle contained a block of 8 no-current trials followed by 8 active-current trials, repeating for a total of 72 trials. The no-current trials allowed us to observe immediate aftereffects (e.g., in launch deviations) following extensive training under the perturbation for the same pair of adapted targets. The cyclical reintroduction of the current after each no-current cycle was intended to maintain the learned compensation for the perturbation and reduce decay between repeated measures of these aftereffects.

#### *Training Phase 2* [150 trials, 75 per Target]

This phase was identical to the Training Phase 1 but instead introduced the novel, transfer target pair. This manipulation allowed us to assess immediate and persistent changes in performance with the novel target pair, following experience compensating for the altered ball-water dynamics during Training Phase 1.

#### *Washout Top-up Phase 2* [72 trials, 36 per Target]

This phase was identical to Washout Top-up Phase 1 but instead presented the second, transfer target pair.

### Trial Termination Criteria and Additional Trial Details

The goal of this task was to launch a virtual ball across a simulated water surface to successfully intersect (i.e., “hit”) a target ring. At the onset of each trial, the target appeared one second after the ball was displayed. This provided participants with sufficient time to perceive the state of the water surface and target location before initiating a throw. If the ball intersected the target, the trial terminated immediately and feedback was presented. For unsuccessful throws, trial termination depended on the water-current state. During no-current trials, the trial ended once the ball had surpassed the target distance, as a hit was no longer possible. During active-current trials, the trial similarly ended once the ball had moved beyond the point at which it could no longer intersect the target under the prevailing water current. This approach prevented unnecessarily long trial durations while preserving all information required to evaluate task performance. Trials also terminated if the ball reached the boundaries of the virtual environment.

At the end of each trial, participants received visual, auditory, and points or reward-based feedback. A line representing the ball’s resulting trajectory was displayed, and a positive or negative tone played depending on the goal state (hit or miss) lasting for one second (see QR code in Figure 2 for a demonstration). Participants also received 5 points when ball hit the target; 3 points if the ball passed within 15.0 cm (four times the target radius); and 0 points otherwise. The auditory feedback and points were meant to encourage participants to consistently aim for the target, while the trajectory line provided participants with knowledge of the movement outcome. After feedback, the next trial began following a one second delay before the subsequent target appeared. The entire task lasted approximately 45 minutes, consisting of a total of 496 trials.

### Procedure

Participants completed a VR ball-launching task where they learned to compensate for a constant leftward acceleration perturbation applied to a projectile’s trajectory. In the virtual environment, the x-axis corresponds to the lateral (left-right) direction, while the z-axis represents the forward (depth). Projectile motion was clamped to the x and z axes; the vertical axis was removed to isolate the effect of the lateral perturbation.

At the onset of each trial, participants were presented with a white ball (diameter = 10.0 cm, virtual mass = 1.0 kg) and one of four spatially distinct targets (diameter= 7.5 cm) positioned at a constant depth, along various positions on the x-axis (See target locations in Figure 2). A single target from a predefined pair was presented in front of the participant along the water surface. To launch the ball, participants first rested the controller near their midline, on a cloth located below their chin (i.e., the ‘home position’). Next, they pressed the ‘trigger’ button using their index finger to initialize the start position while their hand remained at home. To launch the ball, participants then rapidly moved the controller from home along the forward and horizontal directions—with the goal of producing a ball trajectory that passed as close as possible to the target, ideally intersecting it.

Participants were instructed to vary the direction and speed of the controller’s movement to control the ball’s launch trajectory. Controller position was sampled at 72 Hz (matching the headset refresh rate), and the launch vector (used to obtain launch directions and speeds) was computed as the mean velocity of the final 10 samples before exceeding the 30.0 cm displacement threshold, or when the trigger was released. Once the ball was launched, participants received a haptic cue in the form of a controller vibration to indicate the ball has been thrown, prompting them to return the controller back to the home position to observe sensory feedback and prepare for the next trial.

### Water Current Perturbation

A leftward water current was simulated in Unity by incrementally updating the ball’s x-axis velocity on every physics update (Δt = 0.01389 s, 72 Hz). Because this was a novel paradigm, we tested two lateral water current magnitudes in separate experiments: leftward at 2.0 or 3.0 m/s. Given that the ball was 50% submerged when traversing across the water surface, its trajectory was influenced by the simulated water current according to a drag coefficient = 0.47, a fluid density of 1000 kg/*m*^3^, and a ball mass of 1.0 kg. Consequently, the water current produced a gradual arc-like trajectory in the direction of the applied current (see QR code in Figure 2 for a video demonstration). Importantly, the water current acted only on the virtual ball after launch and did not directly alter participants’ movements. The effect of vertical gravity was disabled to isolate adaptation to the lateral water-current perturbation and avoid introducing additional physics interactions to the ball dynamics.

To determine the combination of launch deviation and launch speed required to successfully “hit” each of the target, the task physics were simulated across a range of launch conditions. The resulting solution manifold, defined as the set of execution variable combinations that successfully intersected with a target, is illustrated in Figure 5. Smaller launch deviations required greater launch speeds to maintain task success, whereas larger launch deviations permitted successful performance at lower launch speeds. As illustrated in Figure 5, downstream (leftward) targets afforded a broader range of successful launch parameter combinations than upstream (rightward) targets. Consequently, this creates distinct motor challenges with identical perturbations across the workspace.

**Figure 5:**
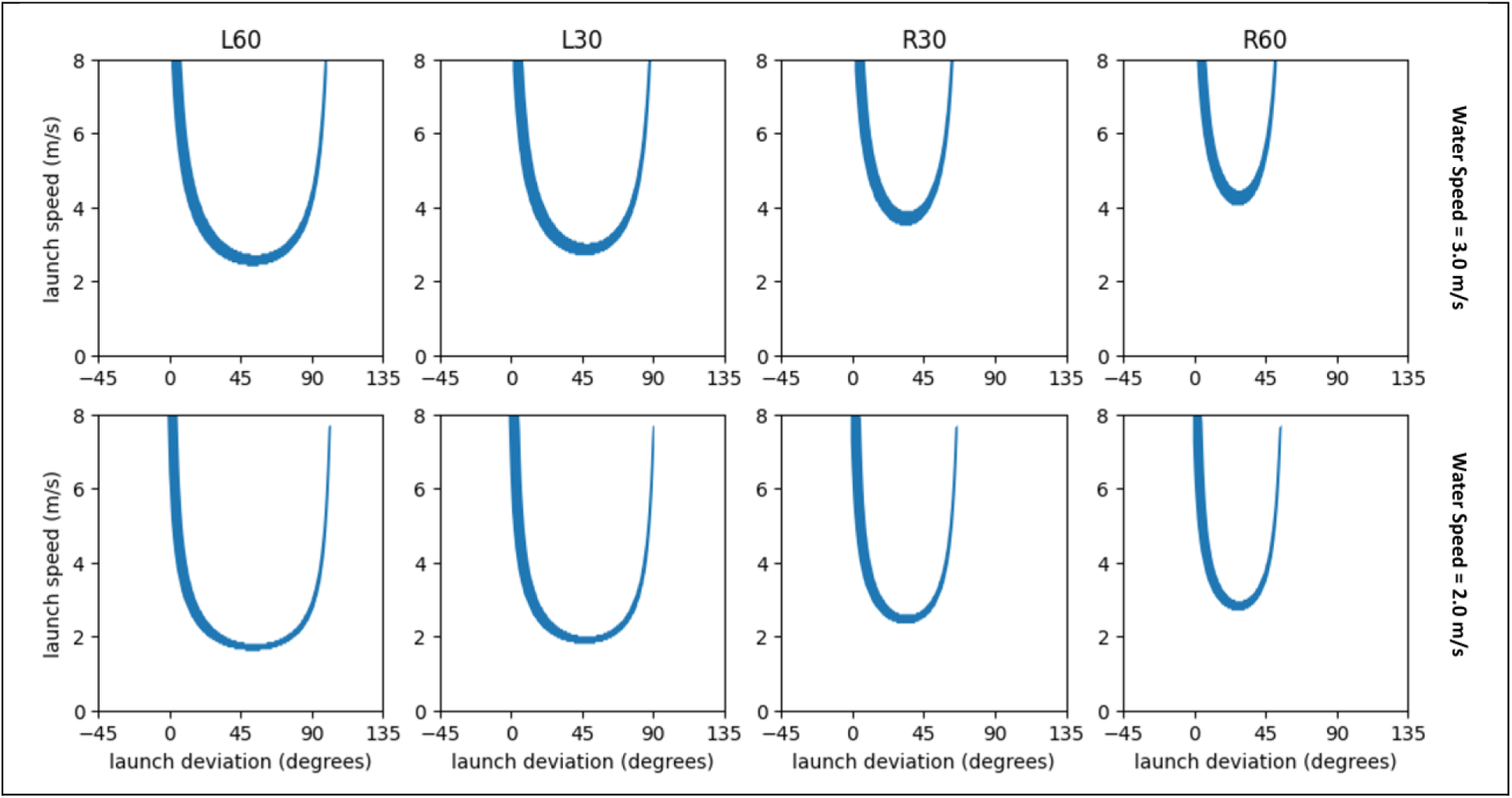
Simulated Solution Manifolds in Deviation and Speed Execution Space. Columns indicate the four target locations (L60, L30, R30, R60), and rows represent the two applied water-current velocities (Top = -3.0 m/s; Bottom = -2.0 m/s). The blue curves illustrate the successful execution variables for each condition. Downstream (Leftward) targets exhibit a wider range of launch angle solutions with lower minimum launch speeds compared to upstream (Rightward) targets. Increasing the water current magnitude from 2.0 m/s to 3.0 m/s shifts the solution manifold vertically along the speed axis, requiring higher launch speeds to hit the target.

### Outcome Measures

To measure task performance, we derived the Euclidean distance between the ball’s (XZ) point of minimum distance and the target’s surface. Because this value was absolute, distances of the same magnitude on either side of the target were indistinguishable. To preserve the signed, relative position of the point of closest approach, we multiplied the Euclidean distance by the sign of the first Cartesian principal component (See Figure 6). To achieve this, we grouped trials by target and water current state conditions and conducted a Principal Component Analysis (PCA) on the X and Z coordinates of the minimum distance points. This allowed us to extract the primary axis of variance for each condition, to determine the signed error (overshoot or undershoot) after the data was projected onto this axis, and to apply the sign of this value back to the absolute Euclidean distance. Furthermore, for all subsequent plots and analyses using this variable, we flipped its sign to reflect an intuitive reduction toward an error of zero.

**Figure 6:**
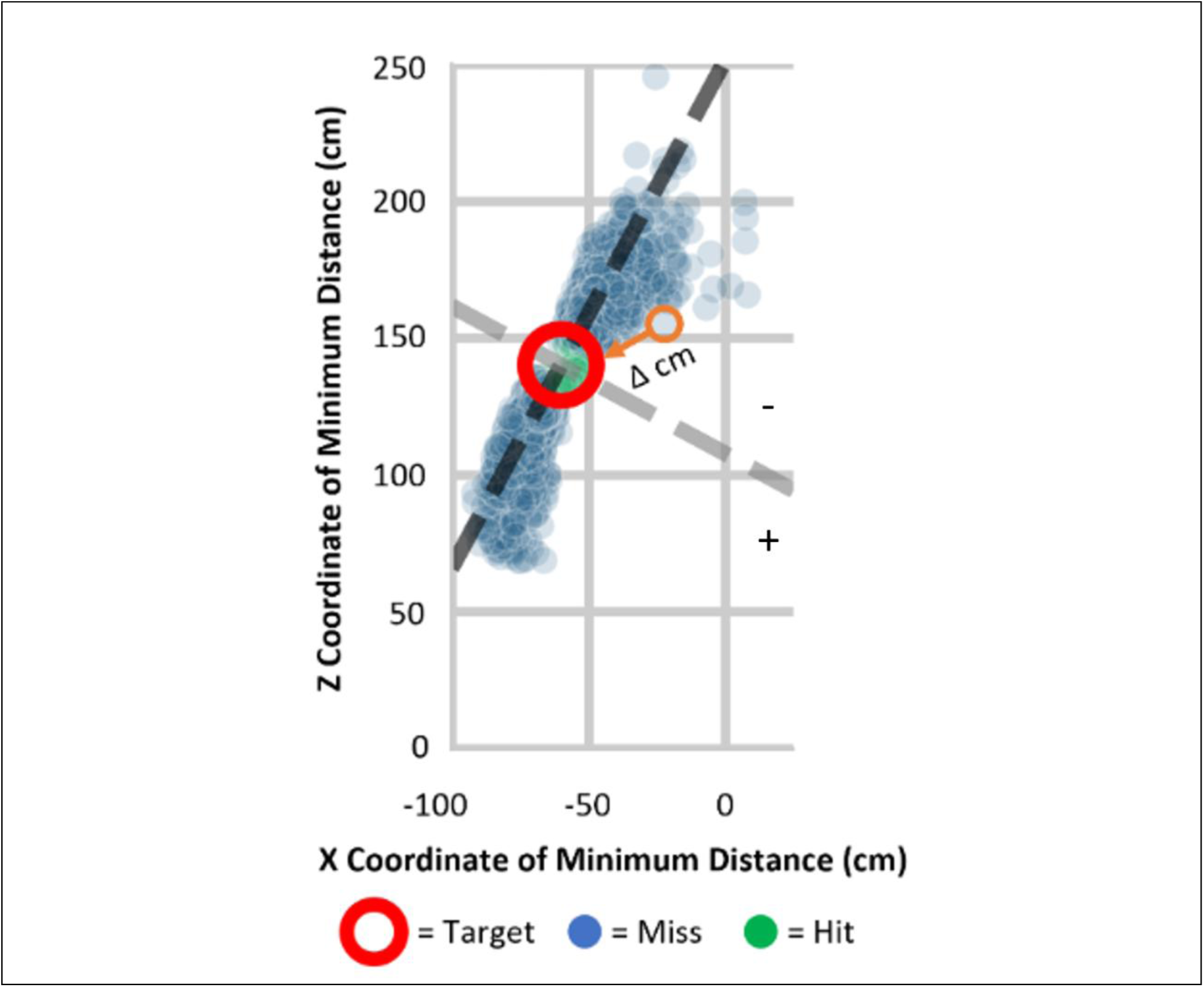
Minimum Metric Error. The sign of the XZ Cartesian position of minimum distance relative to the orthogonal axis (gray dashed line) of the principal component (black dashed line). Minimum Metric Error was derived from the minimum Euclidean distance multiplied by the sign of the XZ position relative to the orthogonal axis. Following the sign flip, positions below the orthogonal axis were assigned as positive, while those above were assigned as negative.

Additional measures of interest were launch deviation (in degrees, relative to target centre) and launch speed (meters/second). Although both variables contributed to task success during active water-current trials, launch deviation was the primary determinant of performance during no-current trials, because successful throws required only that the ball be directed toward the target. Consequently, our aftereffects analyses focused on launch deviation, allowing us to assess whether participants continued to express launch directions appropriate for the previously perturbed environment following current removal. Effective launch deviation hit boundaries spanned ±3.29° and ±3.50° depending on target distance from the initial ball-home position as illustrated by the horizontal bar in Figure 7.

**Figure 7:**
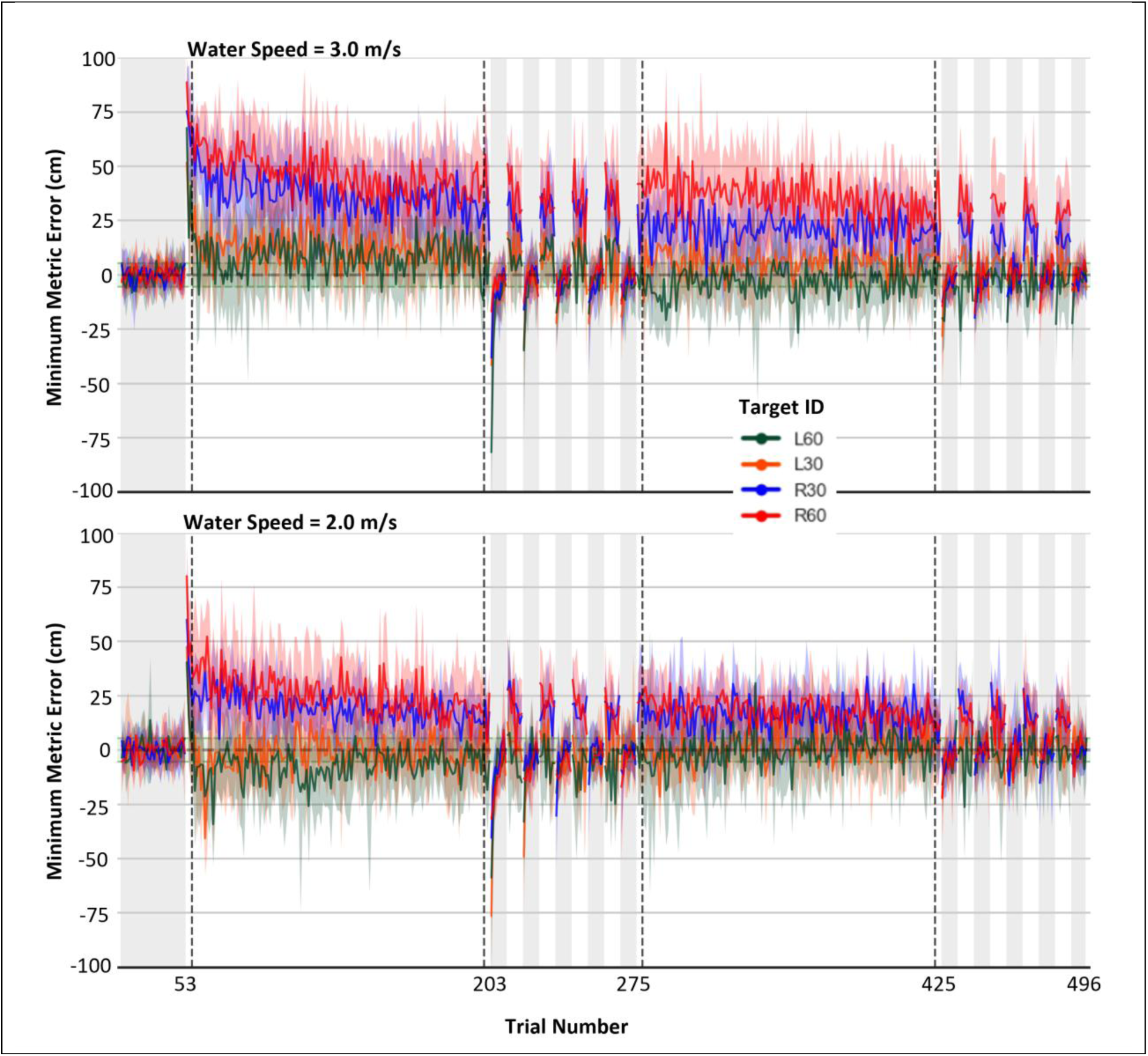
Reductions in Minimum Metric Error across Trials. Each line corresponds to the trial average per target with the 95% confidence interval. Vertically gray shaded trial-spans correspond to no-current trials.

### Data Screening

Participants were excluded if they failed to follow task instructions after clarification and were subsequently replaced for our total of 83 participants. Across all phases, trials were excluded if the ball failed to travel halfway to the target along the z-axis (i.e., 70.0 cm in the forward direction), resulting in the removal of 397 trials (1.02% of all trials). During baseline, trials were excluded if the minimum metric error was greater than 2.0 SD from the Baseline mean, dropping a total of 132 trials (4.97% of all baseline trials). For all subsequent phases, trials were excluded if the minimum metric error exceeded 3.0 SD from each target’s mean minimum metric error, resulting in a removal of a final 62 trials (0.16% of all trials).

### Data Analysis

Because the 2.0 and 3.0 m/s water-current conditions were conducted as separate experiments with separate groups of participants, all analyses were performed separately for each water-current magnitude. For training trials, the primary dependent variable was signed minimum distance to the target (minimum metric error; Figure 6). Minimum metric error was selected because successful performance could be achieved through multiple combinations of launch deviation and launch speed. Consequently, changes in either execution variable alone were not uniquely indicative of learning, whereas minimum metric error provided a direct and consistent measure of task success across the redundant workspace.

To separately evaluate adaptation to each water current, participants were grouped according to the target pair presented during the Training phases. The purpose of this analysis was to establish that participants could reliably reduce performance errors within a single, short training session given the novelty of the task and perturbation. For each water-current experiment, two 2X2 repeated measures ANOVAs were conducted on minimum metric error, with *Trial Set* (early vs. late) and *Target Pair* (L60 and R30 or L30 and R60) as a within-subject factors. Early and late performance were defined as the mean of the first and last two trials of the initial Training Phase, respectively. We chose the first and last two trials to reduce rapid trial-by-trial variability associated with early learning, while still capturing the learning performance immediately following perturbation onset. This analysis was repeated for the second Training Phase to determine whether performance continued to improve following Training Phase 1, or saturated by this learning stage.

To assess transfer, target pairs were counterbalanced across training phases such that participants initially trained on only two of the four targets and encountered the remaining two during the subsequent Training (Transfer) Phase. Consequently, each participant experienced only one target pair within a given phase, precluding a single repeated-measures ANOVA across all targets and phases. Transfer was therefore evaluated separately for each target, directly testing whether performance on the novel transfer target showed clear performance benefits from prior experience with the water-current perturbation. Minimum metric error again served as the dependent variable. For each water-current experiment, four one-tailed Welch’s t-tests (one per target) were conducted with Phase (Training Phase 1 vs. Training Phase 2) as the between-subject factor. One-tailed tests were used because if learning generalized beyond the trained targets, participants encountering a target during Training Phase 2 (experienced state) should exhibit lower initial minimum metric errors than participants encountering the same target during Training Phase 1 (naive state). Thus, a one-tailed test not only aligns with our hypotheses but also provides enhanced power to detect transfer effects. Therefore, transfer was inferred when initial errors for a target were significantly lower during Training Phase 2 than during Training Phase 1. Holm’s correction was applied to account for multiple comparisons.

Aftereffects were evaluated by comparing average launch deviations during Washout no-current trials to the bootstrapped average upper 99% confidence interval (CI) threshold (per target) derived from stabilized baseline and late-training performance (last 8 trials of each phase). Launch deviation served as the primary dependent variable because it was the variable that determined task success during no-current trials. This approach allowed us to identify trials that remained in the previously adapted direction and exceeded stable, baseline behaviour. Namely, launches in the adapted direction (i.e., positive, rightward, greater than 0), and those greater than the baseline average upper 99% CI. We chose the 99% confidence intervals to provide a conservative criterion for identifying aftereffects.

Across all applicable analyses, effect sizes (generalized η² and Cohen’s d) were reported. Importantly, some of the statistical analyses are not fully reported in the text below to make it more readable. However, the full output can be found in the notebook here: [<u>GITHUB LINK</u>]

## Results

### Error Reduction following the first Training Phase

A key signature of motor learning in goal-oriented tasks is the reduction of errors over practice. Adaptation to a perturbation typically manifests as an initial increase in error after perturbation onset, followed by a gradual reduction towards baseline performance. While this is well-established in traditional paradigms such as VMRs, it was important to first establish that participants in our novel environmentally specific perturbation paradigm similarly learned to compensate for the imposed water current within a one hour, VMR-like training period (i.e., approximately one hour). Figure 7 illustrates minimum metric error across the Baseline, Training, and Washout phases for both current magnitudes (2.0 and 3.0 m/s; bottom and top panels, respectively) and targets (different colours), highlighting changes in average performance over time during both current-on (unshaded) and still-water/current-off periods (grey shaded regions).

While Figure 7 illustrates changes in performance outcomes across trials, Figure 8 (A & B) shows the behavioural adjustments (i.e., execution variables: deviation and speed, respectively) associated with these improvements. Figure 8A demonstrates that on average, participants quickly adjusted their launch direction to throw rightward—against the current—when the perturbation was first introduced (Figure 8A, trial 53). However, launch deviation did not remain fixed throughout training and instead showed a modest decline, on average, with continued practice. Similarly, Figure 8B depicts a rapid increase in average launch speed at perturbation onset (Figure 8B, trial 53), followed by a more gradual increase over subsequent training trials. Thus, although participants initially compensated by increasing both launch deviation and launch speed, continued practice was associated with further refinement of these execution variables, reflecting saturation in a lower error region of the execution space.

**Figure 8:**
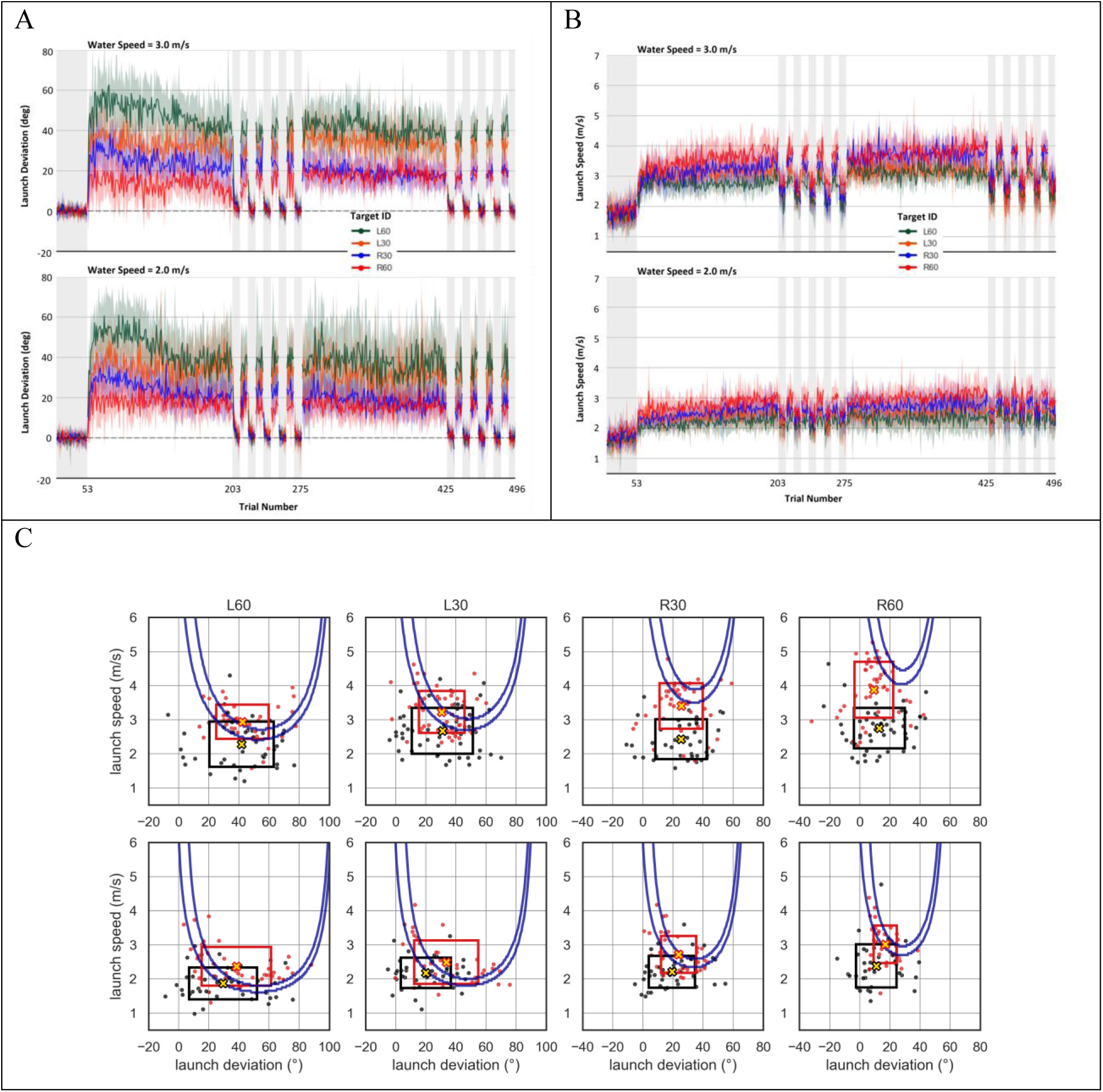
Changes in Launch Deviation (A), and Launch Speed (B) across all Trials in Each Water Speed Experiment. Consistent with Figure 7, each line corresponds to the trial average per target with the 95% confidence interval, while the gray shaded trial-spans correspond to no-current trials. C: Solution manifolds and launch solutions during the first Training Phase, plotted as a function of launch deviation (degrees) and launch speed (m/s). The blue U-shaped regions indicate the combinations of launch deviation and launch speed that resulted in successful task performance for each target. Black and red points represent individual participants’ launch variables for the first and last 2 trials (per target) per participant, respectively. The corresponding black and red boxes illustrate the spread of observed launch variables, with box widths and heights representing variability in launch deviation and launch speed, respectively. Gold ‘x’ symbols indicate the median launch variables associated with each distribution.

While Figures 8A and 8B separately depict changes in launch deviation and speed across all trials, Figure 8C illustrates the solution manifold (blue parabola-like shape) together with executed combinations observed during the first two (black markers) and last two (red markers) trials during Training Phase 1. These trials correspond to the early and late performance measures used in the subsequent statistical analyses. Our data suggests participants primarily shifted the median of the execution variable distributions (as depicted with the gold ‘x’), with a trend illustrating that many targets showed reductions in standard deviations for both execution variables late in Training Phase 1.

To formally assess learning, we compared average minimum metric error between early and late performance within each Training Phase using a series of four 2X2 repeated measures ANOVAs conducted separately for each water current and target pair. Within-subject factors included trial set (early: first two trials, vs late: final two trials) and training target from pair (L60 and R30 or L30 and R60). Figure 7 illustrates the early and late trial-spans with dashed vertical lines during both Training phases.

As illustrated in Figure 9 (Training Phase; left column), we consistently found significant main effects of trial set on minimum metric error for both the 2.0 m/s current (group 1: ; group 2:) and 3.0 m/s current (; group 2:), indicating significant error reduction from early to late training. In all cases, this effect of trial set was consistent across all targets, as target × trial set interactions were not significant following corrections (*p_ajd_* > .09). Therefore, minimum metric error was significantly reduced across all target locations and in both water current experiments during Training Phase 1 (Figure 9 left column). Although performance improved substantially, compensation was not complete for all targets. In particular, errors for the upstream targets (R60 and R30) generally remained elevated compared to baseline performance by the end of training, whereas the downstream targets (L60 and L30) approached baseline levels more closely. Together, these findings demonstrate that participants improved performance in response to the altered ball–water-current dynamics, resulting in robust error reduction following practice; although compensation remained incomplete for some targets.

**Figure 9:**
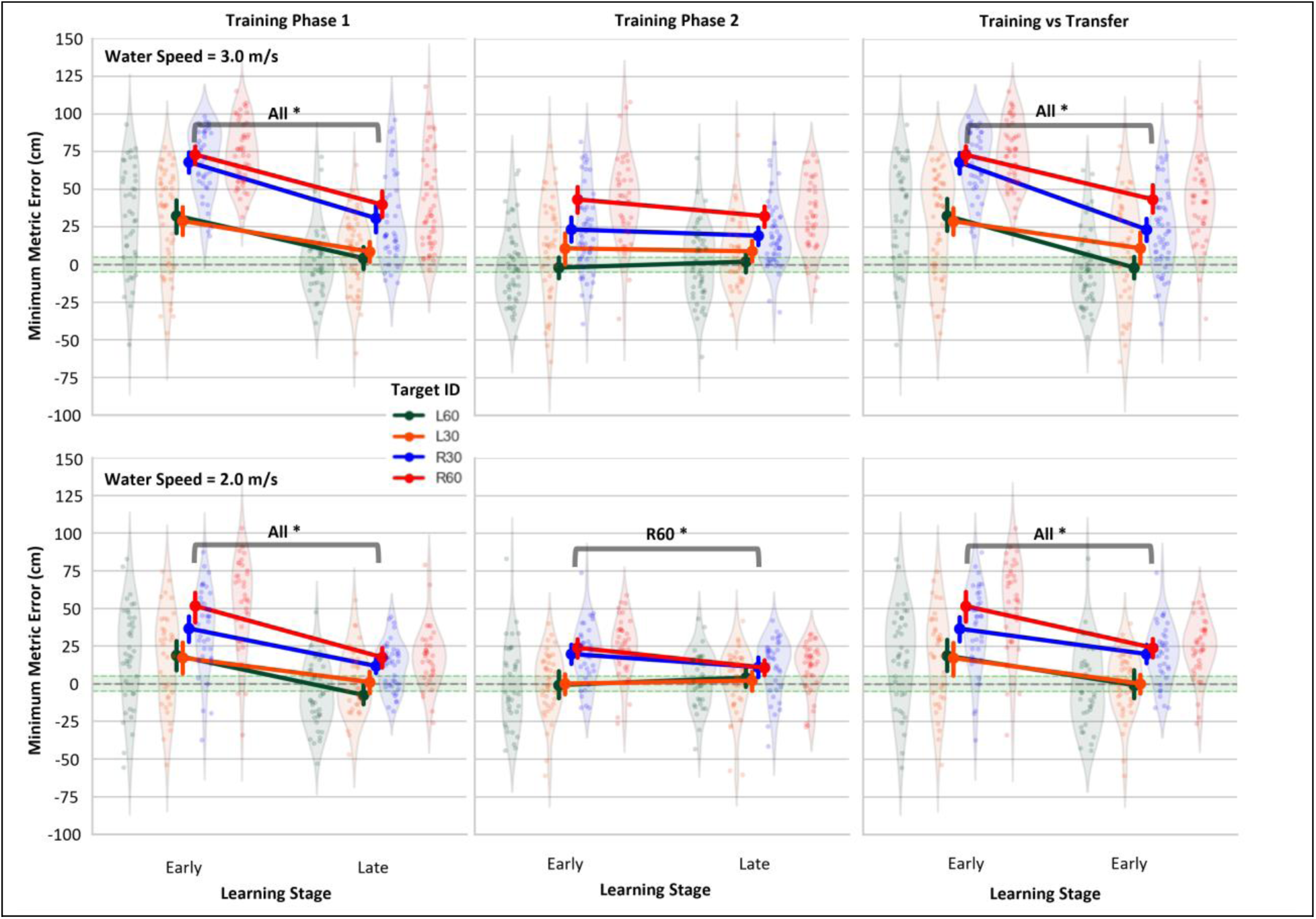
Reductions in Average Minimum Metric Error following Practice. Each line corresponds to the change in average error during key trial sets. Endpoints illustrate the 95% confidence intervals. The horizontal green bands in each facet represents the hit zone: minimum metric errors that overlap the target area and result in successful trials (hits). The violins illustrate individual trials.

For completeness, we also examined whether minimum metric error continued to decrease during the second Training Phase, when the novel target pair was introduced. Using the same analyses described above, we compared early and late performance within this phase (Figure 9, middle column). For the 3.0 m/s current, we did not observe a main effect or interaction with trial set, revealing that learning had peaked or saturated, therefore we did not observe any additional error reductions during this phase (top-middle column). However, for the 2.0 m/s experiment, significant target × trial set interactions were observed for both pair groups (*F* ≥ 5.70, *p* < .05, *η*^2^ ≥ .041) (Figure 9 bottom-middle column). Post-hoc analyses revealed that these effects were driven by the rightward targets, although only R60 remained significant following correction for multiple comparisons. Overall, little additional reductions in minimum metric error were observed in the second Training Phase; with the exception of continued improvement for the R60 target during the 2.0 m/s experiment. These results demonstrate that most of the learning occurred in preceding Training and Washout-Top-Up phases, resulting in learning saturation at the onset of the second Training Phase (Figure 9 Middle). Importantly, because this phase introduced novel targets, the limited additional error reduction observed during the second Training Phase suggests that performance on the new targets may have benefited from transfer of learning acquired earlier in the experiment. We examine this possibility directly in the following section.

### Transferring Learning to Novel Target Locations

Transfer refers to the extent to which learning acquired in one context benefits performance in a novel context. Here, we examined whether adaptation to the water-current perturbation transferred to a second pair of spatially distinct targets following 182 (150 Training and 32 Top-Up) trials under the perturbation. If learning transferred to these novel target locations, participants should exhibit lower initial errors when first encountering the targets during the second Training Phase, than those encountering the same targets during the initial Training Phase. To assess transfer, we compared initial minimum metric errors for each target during Training Phase 1 (i.e., naive state) to initial errors for the same target during Training Phase 2 (i.e., experienced state). Separate one-tailed Welch two-sample t-tests were conducted for each target and water current (2 currents × 4 targets), with Holm’s correction applied to account for multiple comparisons.

#### Speed 3.0 m/s

For all targets, the average initial minimum metric error was significantly reduced for experienced participants during the Training Phase 2 compared to naive participants during the Training Phase 1 (Figure 9 Top-Right). At target L60, naive participants (*M* = 32.46 cm, *SD* = 34.74) demonstrated significantly larger mean initial errors than their experienced counterparts (*M* = -1.87 cm, *SD* = 25.32), *t*(69.24) = 5.23, *p* < .001, yielding a large effect size, *d* = 1.15. A similar pattern was observed for target L30, with naive participants (*M* = 28.91 cm, *SD* = 33.70) yielding larger initial errors than experienced participants (*M* = 10.93 cm, *SD* = 34.41), *t*(82.94) = 2.49, *p* = .007, with a medium effect size, *d* = 0.53. Significant differences were also observed for upstream-rightward targets. For target R30, naive participants (*M* = 68.06 cm, *SD* = 22.30) again showed larger initial errors than experienced participants (*M* = 23.45 cm, *SD* = 28.05), t(88.00) = 8.41, *p* < .001, with a large effect size, *d* = 1.74. Finally, for target R60, naive participants (*M* = 73.03 cm, *SD* = 20.74) showed significantly larger initial errors compared to experienced participants (*M* = 43.42 cm, *SD* = 28.74), *t*(68.78) = 5.48, *p* < .001, and a large effect size, *d* = 1.20.

#### Speed 2.0 m/s

Consistent with the previous results, initial minimum metric error was significantly reduced across all targets for participants during Training Phase 2 compared to naive participants during Training Phase 1 (Figure 9 Bottom-Right). At target L60, naive participants (*M* = 18.72 cm, *SD*= 32.66) showed significantly larger initial errors than experienced participants (*M* = -0.75 cm, *SD* = 28.02), *t*(73.84) = 2.80, *p* < .001, with a medium effect size, *d* = 0.64. Target L30 illustrated a similar pattern with naive participants (*M* = 17.22 cm, *SD* = 31.63) demonstrating a significantly larger initial error than experienced participants (*M* = 0.19 cm, *SD* = 21.48), *t*(60.70) = 2.72, *p* < .001, and a medium effect size, *d* = 0.64. Again, the upstream-rightward targets demonstrated a similar effect. For target R30, naive participants (*M* = 36.68 cm*, SD* = 27.43) showed larger initial errors than experienced participants (*M* = 19.87 cm, *SD* = 20.10), *t*(71.19) = 3.07, *p* < .001, with a medium effect size, *d* = 0.69. Lastly, for target R60, naive participants (*M* = 51.69, *SD* = 31.05) had larger initial errors than experienced participants (*M* = 24.00, *SD* = 19.62), *t*(57.97) = 4.59, *p* < .001, and a large effect size*, d* = 1.08.

Taken together, these results demonstrate that experience with the perturbation yields immediate transfer to novel spatial targets, indicating strong transfer of the learned, compensatory motor commands.

### Motor Aftereffects during Washout

A signature of motor adaptation is the emergence of aftereffects: persistent-compensatory movements following the removal of the perturbation. Thus, after establishing robust error reduction and transfer, next we investigated whether aftereffects were evident during the no-current trials of the Washout phases (see Figure 10).

**Figure 10:**
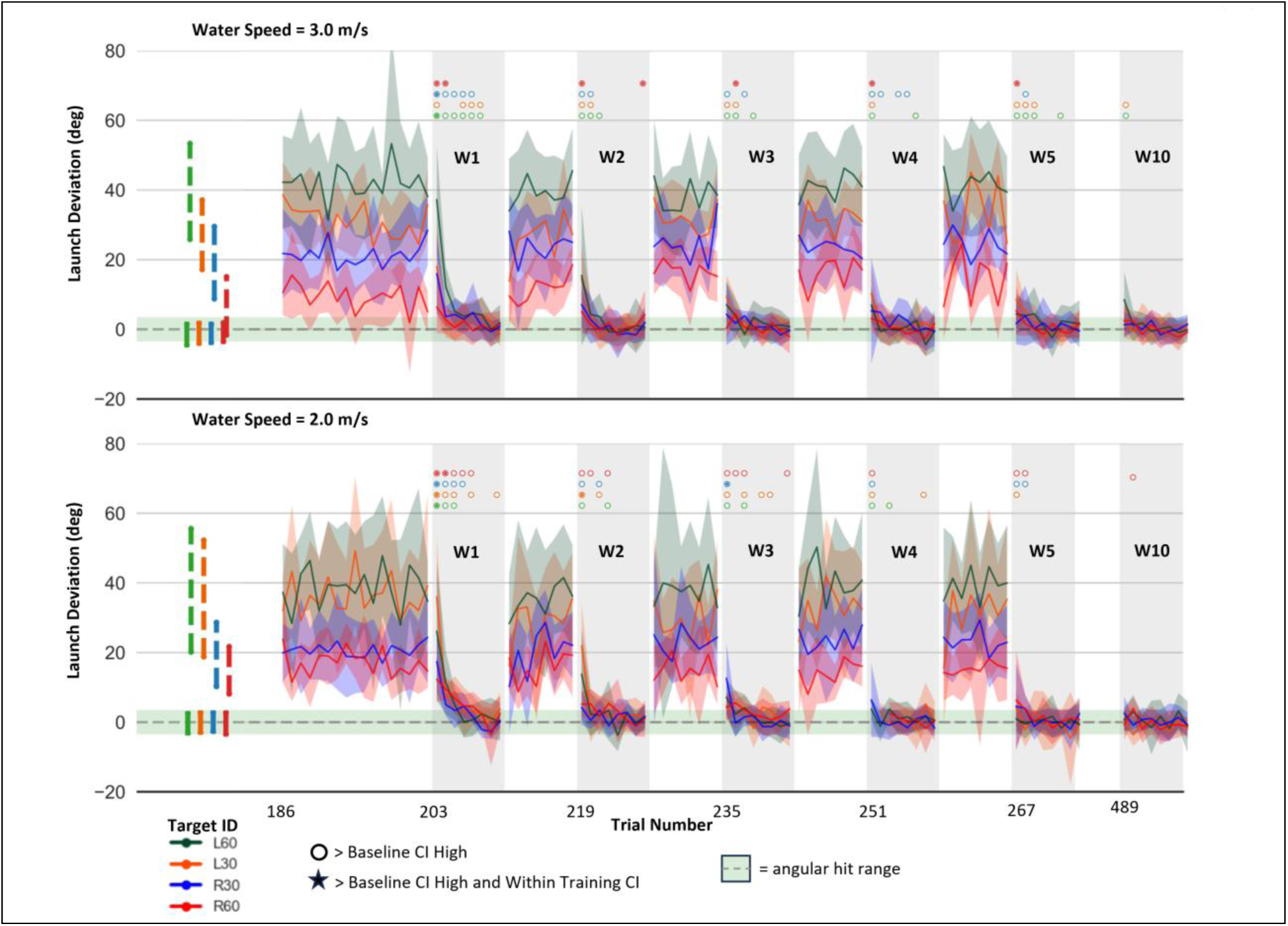
Aftereffect Frequency. Zoomed in Figure 8A, illustrating the final 16 Training Phase 1 trials, Washout-Top-Up Phase 1 (W1 – W5), and the final no-current cycle of Washout-Top-Up Phase 2 (W10). Gray, vertically shaded trial-spans depict the no-current Washout trials interspersed between active-current trials. The solid coloured vertical lines centred around zero depict the 99% Bootstrapped CI during the final eight trials (per target) during Baseline. Additionally, the horizontal green span reflects the widest, successful launch deviations during washout (i.e., ±3.50° for ±30.0 cm targets; ±3.29° for ±60.0 cm targets). The dashed-coloured vertical lines above the Baseline CIs illustrate the 99% Bootstrapped CI during the final eight Training Phase 1 trials (per target).

Here, we focused our analysis on changes in launch deviation relative to the target, as it was the primary determinant of task success during Washout no-current trials. Because aftereffects were expected to be largest immediately following each context transition and diminish rapidly thereafter, we examined their presence by comparing trial-by-trial performance against baseline and training 99% confidence intervals; rather than conducting separate statistical tests for each Washout trial.

As illustrated in Figure 10, average launch deviation for nearly all targets in both experiments fell outside the baseline 99% CI during the first trial in Washout Cycle 1 (W1), indicating robust aftereffects immediately following perturbation removal (circles, in Figure 10). Furthermore, nearly all targets (except L30 in the 3.0 m/s experiment) exhibited average deviations that were also within the 99% CI during the final 8 trials of the Training Phase (stars-in-circles, in Figure 10), indicating that participants were consistently executing launch directions appropriate for the prior perturbed context. The magnitude of the first-trial aftereffect varied across targets, and closely mirrored the target-specific deviation gradient observed at the end of training (See Figure 10, before trial 203). Thus, targets requiring larger compensatory launch deviations during training generally exhibited larger aftereffects following perturbation removal. Importantly, these aftereffects were observed despite a one-second delay before target presentation and the absence of the audiovisual cues associated with the perturbation.

Although launch deviations decreased rapidly following the initial transition to no-current trials, many launches for subsequent trials continued to exceed the baseline 99% CI, indicating persistent aftereffects beyond the first Washout trial (as indicated by the hollow circles). In many cases, aftereffects remained evident for 2-3 additional, consecutive no-current trials and occasionally as many as five. Across subsequent Washout cycles, numerous trials continued to exceed the baseline 99% CI (Figure 10, circles), although the frequency of these occurrences progressively declined. This reduction in the occurrence of aftereffects suggests that participants became increasingly effective at adjusting behaviour across repeated transitions between current-on and current-off contexts. Several early Washout trials also remained within the training 99% CI (Figure 10, stars), indicating aftereffects comparable in magnitude to the compensatory deviations expressed during training. This may suggest that participants were unaware of the context change until receiving unperturbed trajectory feedback. This pattern continued across later Washout no-current trials, but we observed a declining trend in aftereffect frequency with each cycle, with almost no aftereffects in the final Washout Phase 2 cycle (See Figure 10, W10).

## Discussion

Motor learning is often studied using perturbations that directly alter limb movements or the relationship between movements and sensory feedback. In the present study, we examined whether key signatures of adaptation would extend to a task in which a visible environment-force acted exclusively on a manipulated object after release, and whether successful performance could be achieved though multiple combinations of launch direction and speed. Participants exhibited robust error reduction to all targets following training, indicating substantial adaptation to the altered ball–water dynamics, generalized this learning to novel target locations, and expressed persistent aftereffects following perturbation removal. Together, these findings demonstrate that humans can acquire predictive action–outcome mappings—an internal model or representation—of object–environment dynamics under redundant task conditions, and that many of the classic signatures of sensorimotor adaptation extended beyond traditional limb-based perturbations.

### Participants reduced errors under the water-current perturbation

In any sensorimotor adaptation paradigm, the CNS must recalibrate a predictive mapping between motor commands and sensory expectations to account for altered environmental dynamics. In goal-oriented tasks, this adaptation typically appears as a gradual reduction in performance error as internal model expectations are updated to compensate for the imposed perturbation. In the case of traditional limb-based perturbations such as VMRs, sensory discrepancies prompt the CNS to continuously update the estimated state of the manipulated effector until no discrepancies remain (Tseng et al., 2007). For tasks like VMRs, successful performance can be achieved by adjusting a single execution variable and applying the adjustment across the workspace. In contrast, successful performance in the present task required coordinating changes in at least two execution variables—launch direction and speed—dynamically across the workspace. Here, adaptation is not the result of a simple scalar adjustment of one execution variable, but rather from altering and covarying the manipulation of multiple variables to maintain task success (Zhang et al., 2018; Zhang & Sternad, 2021). Participants in our task nonetheless showed robust reductions in performance error across all targets and both water-current magnitudes, demonstrating successful adaptation to the altered ball–water dynamics. Therefore, the consistent error reduction observed across the workspace demonstrates that participants developed an experienced-driven, predictive mapping between launch parameters and resulting ball trajectory.

While Figure 7 illustrates reductions in performance error, Figure 8 provides additional context regarding the behavioural changes associated with this learning. Figures 8A and 8B reveal that on average, participants initially compensated by increasing both launch deviation and speed following perturbation onset. With continued practice, these execution variables continued to evolve, suggesting that participants refined their launch behaviour beyond their initial compensation. Although these changes are descriptive rather than formally quantified, they are consistent with participants learning to exploit multiple execution variable combinations to improve task performance, as demonstrated in Figure 7.

Unlike traditional VMR tasks, participants could not compensate by applying a simple directional correction. Instead, successful performance required anticipating how the water current would influence the ball’s trajectory after release and coordinating both launch direction and launch speed accordingly. Thus, adaptation extended beyond recalibrating the relationship between limb movement and sensory feedback, to developing a predictive mapping of how an environmental force will impact a manipulated object following a self-generated ballistic movement.

Interestingly, adaptation was not uniform across targets. In general, targets associated with narrower solution manifolds exhibited greater residual error following training than targets with broader solution manifolds. Likewise, we observed ongoing significant error reduction for the R60 target during Training Phase 2 under the 2.0 m/s current. This suggests that the geometry of the solution manifold may influence the ease with which participants identify and consistently engage successful launch solutions. However, this pattern was not universal, as the leftward targets in the 2.0 m/s condition showed comparable adaptation despite its broader solution manifolds. One possible explanation is that narrower solution manifolds require greater precision in coordinating execution variables because fewer combinations produce successful outcomes. Additionally, because these narrow manifolds require higher minimum speeds for success (see Figure 5), success likely introduces additional motor variability through signal dependent noise associated with increasingly vigorous movements. Future studies that systematically manipulate solution-manifold geometry independently of perturbation magnitude will be required to directly test this possibility.

Recent work from our and other labs has shown that the CNS can adapt to a variety of perturbations applied to manipulated virtual objects after movement completion—including imposed rotations, curves, accelerations, and altered gravity (King et al., 2026; Modchalingam et al., 2025; Pouke et al., 2024). The present findings extend this growing body of work by demonstrating successful adaptation to a continuous environmental force acting on a manipulated object within a redundant execution space. An important remaining question, however, is whether this learning reflected compensation specific to the trained targets or a more general representation of the underlying ball–environment dynamics. We address this question by examining transfer to novel target locations below.

### Learning transferred strongly to novel target locations

A defining characteristic of motor adaptation is that learning generalizes beyond the specific movements experienced during training. If participants had simply memorized target-specific launch solutions during Training Phase 1, switching to the novel targets would be expected to produce substantial performance errors and require renewed adaptation. Instead, participants demonstrated substantial, immediate transfer to the novel target pair, with performance at the beginning of Training Phase 2 significantly different from performance at the start of Training Phase 1. Furthermore, continued practice on the transfer targets produced little additional improvement, indicating that most of the learning acquired during the initial training phase generalized to the novel target locations.

This effect was consistent across both water speed experiments and all targets, where experienced participants demonstrated significantly lower initial performance errors compared to naive participants exposed to the same targets for the first time. Importantly, each target was associated with a distinct solution manifold, requiring different combinations of execution variables to achieve task success. Because transfer was observed despite distinct solution manifolds, this adaptation cannot be from a simple memorization of successful target-specific execution variables. Rather, participants appeared to have acquired knowledge that generalized across target locations within the same environmental context. This predictive action–outcome mapping of ball-water dynamics facilitated immediate transfer benefits to novel targets, despite featuring different solution manifolds compared to previously trained targets.

Our transfer results may be enhanced by the degree of movement similarity shared by adjacent targets between phases. Transfer in traditional VMR paradigms has been explained by overlapping neural tuning curves centred on the trained or intended movement direction, where adaptation maximally generalizes to adjacent locations within 45 degrees (Day et al., 2016; Krakauer et al., 2000; Tanaka et al., 2009). Therefore, our transfer results are consistent with previous work suggesting that adaptation generalizes strongly to similar movement directions.

Another key factor that may contribute to generalization within this task is how the CNS infers the source of errors. Because state uncertainty for the body, tool, and environment is omnipresent, the CNS uses sensory observations to better infer the state of these error sources to update the appropriate internal model (Berniker & Körding, 2008; Wei & Körding, 2009). Importantly, how the CNS infers these states determines not only which internal model is updated, but also how this learning generalizes.

The present task was designed to reduce this state uncertainty by introducing explicit sensory cues the CNS can use to infer the state of the external environment. Specifically, the water current perturbation was associated with visual motion of the water surface, the sound of flowing water, and the perturbed trajectory of the launched ball. Together, these cues likely indicated to the participants that the external environment is causing errors and thus contribute to updating an internal model of the environment’s impact on the ball, rather than an internal model of the effector. Because these errors are primarily caused by external sources and not entirely by the effector, we would expect the resulting generalization curve to be broad and transfer across limbs (Berniker & Körding, 2008). Although the present study did not directly manipulate state uncertainty, future work could examine whether reducing or altering the reliability of these environmental cues influences the extent to which learning generalizes to novel targets.

Because the novel targets contained shifted solution manifolds, memorizing local launch solutions and simply applying them to the novel targets would not have resulted in successful performance. Instead, participants exhibited strong immediate transfer suggesting they learned a context-dependent, predictive action-outcome mapping that generalized to new target locations requiring different launch solutions. Therefore, our results suggest participants developed a generalizable internal model of object-environment dynamics rather than collection of target-specific launch solutions.

### Aftereffects persisted despite visible removal of the perturbation and other relevant cues

While the transfer results provide evidence that participants developed a generalizable internal model of object-environment dynamics, the presence of persistent motor aftereffects highlights the predictive nature of this model. Behaviorally, motor aftereffects appear as compensatory actions even after the perturbation is removed, reflecting a predictive mapping of actions and sensory consequences (Shadmehr & Mussa-Ivaldi, 1994; Wolpert et al., 2011). Here, we manipulated the presence of the water current to assess motor aftereffects. Figure 10 demonstrates large compensatory aftereffects during early Washout cycles (W1 to W5), where launch deviations remained outside the Baseline 99% CIs for many consecutive trials. This demonstrates that, on average, participants initially executed motor commands consistent with the expectation of a perturbed ball trajectory. Importantly, while many traditional visuomotor tasks provide limited cues to indicate perturbation removal—besides sudden unperturbed visual feedback—the present task removed visual water motion and audio cues associated with the perturbation, then presented the target after a 1.0 second delay, and even displayed an unperturbed trajectory line as knowledge of results. Interestingly, despite these explicit cues and the delay in target presentation, aftereffects still emerged.

This effect appeared relatively consistently across the workspace and in both experiments (see associated markers in Figure 10). Motor aftereffects like these suggest participants anticipated a perturbed trajectory to follow the executed motor command, thereby generating a sensory prediction error (SPE). This provides further evidence of a predictive mapping of object-environment dynamics because participants expected the water current to alter the ball trajectory, prompting the expression of adapted motor commands. Furthermore, the fact that many subsequent trials exhibited launch deviations beyond baseline—despite conscious awareness of the context change provided from clear cues and trajectory feedback—suggests that this model was not easily washed out. However, as illustrated in the subsequent Washout cycles, there is a noticeable decreasing trend in the frequency of aftereffects, with the fewest in the final Washout cycle (Figure 10). Rather than an immediate context-switch at the onset of Washout, this decreasing trend of aftereffects within and across cycles demonstrates that participants were becoming more efficient at contextual inference and memory retrieval (Heald et al., 2021).

While participants did not receive online visual feedback of their aiming direction before the ball launched, one could argue that performance in this task relies partially on conscious aiming strategies. This viewpoint is supported by the reductions in launch deviation after trial 1 of the first Washout cycle. Here, upon observing the water current had been removed, participants rapidly reduced launch deviations toward zero. However, because compensatory aftereffects persisted beyond baseline behaviour for several additional Washout trials, explicit strategies alone are unlikely to fully explain these observations. Instead, this gradual deadaptation is consistent with a continued contribution of implicit processes driven by SPEs. However, because the task was not designed to distinguish between explicit and implicit mechanisms, future studies will need to separate the relative contributions of these processes.

Overall, participants continued to generate motor commands consistent with expectations of a perturbed trajectory, even after clearly removing cues associated with the perturbation. Together, the presence of generalization and aftereffects points to the development of a predictive action-outcome mapping, and thus, an internal model of object-environment dynamics, rather than the simple trial-by-trial refinement of strategic motor commands.

### Explicit, implicit, and reward-related contributions to learning

One feature of the present paradigm is that participants received multiple learning signals, including visual trajectory feedback, auditory outcome feedback, and rewards based on task success. While these elements were intended to maintain engagement and provide clear knowledge of results, the present design does not allow us to determine the relative contribution of sensory prediction errors, explicit strategic adjustments, and reward-based learning processes to adaptation (Taylor et al., 2014; Tsay et al., 2022). Future work could independently manipulate trajectory feedback, reward, and environmental cues to determine their relative contributions to learning and transfer in environmentally specified perturbations.

### Limitations and Future Directions

While this novel virtual reality paradigm provides rich insights into motor learning under environmental perturbations, there are a few methodological limitations to consider. First, targets were arranged along a constant depth axis to ensure they fall along a consistent axis parallel to the lateral water current. While this was done to standardize the physics applied to the ball when attempting to hit each target, it resulted in small differences in the Euclidean distance from the origin between the inner (L30 & R30) and outer (L60 & R60) targets. Although this distance difference was marginal, it does introduce varying visible spans and constrained 1D solution manifolds during no-current trials. Future studies using this paradigm could arrange the targets radially to ensure equivalent Euclidean distances and visible angular spans across targets.

Like Modchalingam et al. (2025), there were two ways to release the ball—by moving the controller beyond a predefined distance threshold after an initial trigger press, or to simply release the trigger during the movement. This, however, may introduce an additional execution variable to be manipulated for task success (i.e., release timing) which can be used to modulate initial release velocity. Future studies should either standardize the release method or explicitly measure release timing to determine whether it contributes to performance.

Finally, adaptation was also incomplete for some targets, particularly the rightward targets, and to a larger extent under the 3.0 m/s current. While the exact cause of incomplete adaptation remains unclear, potential explanations include interference from practicing with two targets, or perhaps full compensation simply requires more trials.

To address these possibilities, future studies could determine if adaptation reaches an asymptote and whether that asymptote reflects full compensation after extended practice. Additionally, subsequent work could investigate if learning differs when training on one versus two targets, or how learning evolves across multiple days of practice.

## Conclusion

This thesis aimed to characterize sensorimotor adaptation to altered object-environment dynamics using a novel virtual reality ball-launching task in which participants learnt to compensate for a simulated water-current perturbation. Consistent with classic findings from visuomotor and force-field adaptation paradigms, participants demonstrated key signatures of sensorimotor adaptation—namely, error reduction during training, substantial transfer to novel target locations, and persistent aftereffects following perturbation removal. Together, these findings indicate that humans can rapidly acquire and generalize predictive-mappings between their actions and resulting object motion under altered object-environment dynamics, even within a redundant solution manifold that affords multiple successful launch strategies. More broadly, the present paradigm bridges classical sensorimotor adaptation with the control of manipulated objects in dynamic environments, providing a foundation for investigating how humans learn, generalize, and flexibly update predictive motor control in more naturalistic settings.

